# Modulation of TOR kinase activity in *Chlamydomonas reinhardtii*: Effect of N-starvation and changing carbon pool

**DOI:** 10.1101/312942

**Authors:** Shivani Upadhyaya, Shreya Agrawal, Anmol Gorakshakar, Basuthkar Jagadeeshwar Rao

**Affiliations:** Department of Biological Sciences, Tata Institute of Fundamental Research (TIFR), Homi Bhabha Road, Colaba, Mumbai – 400005, INDIA.; Max Planck Institute of Molecular Plant Physiology, Potsdam - 14476, Germany.; School of Biosciences and Technology, VIT University, Vellore, Tamil Nadu – 632014, INDIA.

**Keywords:** TOR kinase activity, *Chlamydomonas* S6 Kinase, Glucose, Autophagy, Lipid accumulation, N starvation, Hexokinase1

## Abstract

<u>**T**</u>arget <u>**O**</u>f <u>**R**</u>apamycin (TOR) kinase is a sensor as well as a central integrator of internal and external metabolic cues. The upstream and downstream signals of this kinase are very well characterized in animals. However, in algae, higher plants and other photosynthetic organisms, the components of the TOR kinase-signaling are yet to be characterized. Here, we establish an assay system to study TOR kinase activity in *C.reinhardtii* using the phosphorylation status of its downstream target, CrS6K. We further use this assay to monitor TOR kinase activity under various physiological states such as photoautotrophy, heterotrophy, mixotrophy and nitrogen starvation. We observe that autotrophy in light (and not in dark) leads to TOR kinase attenuation during N starvation while the same is not observed in mixotrophy. Importantly, we show that the external carbon source glucose is sensed and uptaken by *C.reinhardtii* cells only in the presence of light and not in the dark. And such exogenously added glucose, as the photoassimilate carbon mimic, results in enhanced production of ROS, induction of autophagy and concomitant drop in TOR kinase activity, creating N-starvation-like cellular state even in N+ conditions. Interestingly, dose dependent addition of glucose revealed TOR kinase activation in low glucose regime (ROS independent) followed by attenuation of TOR kinase (ROS dependent) at high glucose levels.

**Summary:** TOR kinase activity in *C.reinhardtii* is modulated by available carbon source especially glucose, where low levels of glucose cause an increase whereas high levels cause a reduction in TOR kinase activity.

## INTRODUCTION

One of the key regulators of cellular metabolism and growth is the TOR (Target of Rapamycin) kinase. In mammals and yeast, it is known that TOR kinase senses and responds to amino acids, sugars, ATP levels and many other metabolically important signals (Gonzalez *et al.*, 2017; Loewith *et al.*, 2011; Sabatini, 2017). TOR kinase activity and regulation including its upstream and downstream targets in *C.reinhardtii* or other photosynthetic organisms are not fully understood (Perez-Perez, Couso, & Crespo, 2017; Schepetilnikov *et al.*, 2018). However, due to recently developed endogenous assay system to monitor TOR kinase activity in *Arabidopsis*, many new components in TOR signaling, unique to photosynthetic organisms, have been identified. There are recent reports that the plant TOR kinase has evolved to sense and respond to signals unique to plants such as photoassimilate sugars, plant phytohormones auxin, brassinosteroids and abscissic acid receptors (Li *et al.*, 2017; P. Wang *et al.*, 2018; Xiong *et al.*, 2013; Zhang *et al.*, 2016). A recent study has also deciphered the mechanism of regulation of TOR kinase by auxins where the light induced auxin signal in *Arabidopsis* is transduced to TOR kinase by a small GTPase Rho-related protein 2 (ROP2) (Li *et al.*, 2017).

In *C.reinhardtii*, studies on TOR kinase have revealed its importance in regulating autophagy, ER stress and oxidative stress (Diaz-Troya *et al.*, 2011; Perez-Martin *et al.*, 2014; Perez-Perez *et al.*, 2012; Perez-Perez, Couso, & Crespo, 2017). Recent studies in this alga using the classical TOR kinase inhibitor rapamycin have demonstrated the role of TOR kinase in maintaining the Carbon/Nitrogen balance (Juppner *et al.*, 2018). The role of TOR kinase in cellular phosphate metabolism was also uncovered in TOR hypersensitive mutants revealing lower levels of InsP_6_ and InsP_7_ (Inositol Phosphates) (Couso *et al.*, 2016). However, due to lack of an endogenous assay to monitor TOR kinase activity in *C.reinhardtii*, mechanistic understanding of the upstream and downstream components of TOR kinase pathway has remained unclear.

The major advantage of *C.reinhardtii* as an experimental system is that it can be grown in multiple metabolic states, which include photoautotrophic (no organic carbon source provided - grown in light), heterotrophic (carbon source provided – grown in dark) and mixotrophic (carbon source provided – grown in light) modes, involving different regulatory mechanisms (E. H. Harris, 2001; Stern, 2009). All these physiological states impart certain level of metabolic plasticity to the algae, but how these states regulate growth and metabolism via TOR kinase is not well understood. The regulation of these states by TOR kinase might also help uncover insights on how TOR kinase activity is modulated by these states.

To probe this aspect, we begin by establishing an assay system to study TOR kinase activity using pCrS6K (phosphorylated *C.reinhardtii* S6 Kinase) as its functional readout. We observe that, during nitrogen starvation, TOR kinase activity is attenuated in the presence of light while remaining unchanged in dark. This suggests that light or photosynthetic output in the form of fixed carbon could play an important role in regulating TOR kinase activity. Upon providing exogenous glucose as the photoassimilate carbon under photosynthesis inhibition by atrazine or N starvation, we observed an attenuation of TOR kinase activity indicating the role of carbon source glucose in negatively regulating the TOR kinase activity. However, it has been reported that exogenous glucose is not uptaken in dark (Doebbe *et al.*, 2007), whereas the uptake of glucose in light is not well understood. Our studies with fluorescently tagged deoxyglucose in light versus dark indicated that exogenous glucose is uptaken by *C.reinhardtii* only in the presence of light and that high concentration of glucose results in attenuation of TOR kinase activity. We further showed that this exogenous glucose results in ROS (Reactive Oxygen Species) accumulation in the cells that results in induction of autophagy. Finally, we show that glucose mediated change in TOR kinase activity follows a bi-phasic response: first phase where TOR kinase activity rise is perhaps ROS independent and glucose (or glucose derived metabolites) dependent followed by the second phase where increase in ROS leads to attenuation of TOR kinase activity at high glucose level. We discuss this model further integrating other results described here.

## RESULTS

### Identification of a putative *C.reinhardtii* S6 Kinase (CrS6K)

S6K (Ribosomal protein S6 kinase) is one of the best characterized downstream targets of TOR (**T**arget **O**f **R**apamycin) kinase (Saxton *et al.*, 2017). It is a substrate of TOR kinase in human, yeast and *Arabidopsis* TOR-S6K pathway and is known to promote protein synthesis. The same in *C.reinhardtii* pathway is yet to be characterized as reiterated in the earlier reports (Ma *et al.*, 2009; Perez-Perez, Couso, & Crespo, 2017; Xiong *et al.*, 2012). The multiple sequence alignment using MUSCLE (<u>**MU**</u>ltiple <u>**S**</u>equence <u>**C**</u>omparison by <u>**L**</u>og-<u>**E**</u>xpectation) (Edgar, 2004) revealed the existence of a sole S6K gene in *C.reinhardtii* annotated genome with 35% and 40% putative protein sequence identity with *Arabidopsis* and *Homo sapiens* S6K1 respectively, where the C-terminal region was highly conserved. The *C.reinhardtii* genome version v4.0 (https://genome.jgi.doe.gov/Chlre4/Chlre4.home.html) showed that S6K1 is a single gene with the coding sequence length of 3006 bp, constituting 7 introns and 8 exons, yielding a putative protein with the annotated molecular weight of 100KDa(https://phytozome.jgi.doe.gov/pz/portal.html#!gene?search=1&detail=1&method=4614&searchText=transcriptid:30784665). Furthermore, bioinformatic analysis revealed that the N-terminal region of the protein is highly divergent which consists of repeats of amino acids such as poly A, G, Q and poly P as depicted in red (Fig. 1A). When we tried to identify the conserved domains in the protein using the Conserved Domain Database (CDD) search service of NCBI (https://www.ncbi.nlm.nih.gov/Structure/cdd/wrpsb.cgi), we found that the long stretch of amino acids towards the N-terminal did not correspond to any known conserved protein domain. This suggests that the N-terminal region of S6K1 protein in *C.reinhardtii* probably does not have any defined structural or functional domains and hence could be spliced off in the final mature coding transcript. Thus, it is likely that the *C.reinhardtii* S6K is a much shorter polypeptide as in *Arabidopsis* (Xiong *et al.*, 2012) and human S6K1 (Tavares *et al.*, 2015) that also show shorter versions of 46.5 and 70 KDa molecular sizes, respectively. The UniProt database also suggests the S6K1 molecular weight of 35.5 KDa (http://www.uniprot.org/uniprot/A8HTN0) in *C.reinhardtii,* a size shorter than the annotated full-length polypeptide. Moreover, Its C-terminal conserved domain retains the TOR kinase-phosphorylating target motif FLGFTYVAP (Schalm *et al.*, 2002) with the Threonine residue as the TOR phosphorylation target (Colored green in Fig. 1A). The entire TOR phosphorylating motif (shown in the box, Fig. 1A) is conserved in *C.reinhardtii* except for one residue at −3 position (counted with respect to the conserved Thr residue), which is Asp in *C.reinhardtii* instead of Leu in human. With this information, we went ahead with detecting the putative *C.reinhardtii* S6K (CrS6K) protein using a commercial antibody that detects bulk human S6K protein (70 KDa) (Cell Signaling Technology Cat no - 9202S) whose epitope is in the conserved C-terminal region of the protein and hence is expected to cross-react with *C.reinhardtii* S6K1 too.

**Figure 1.**
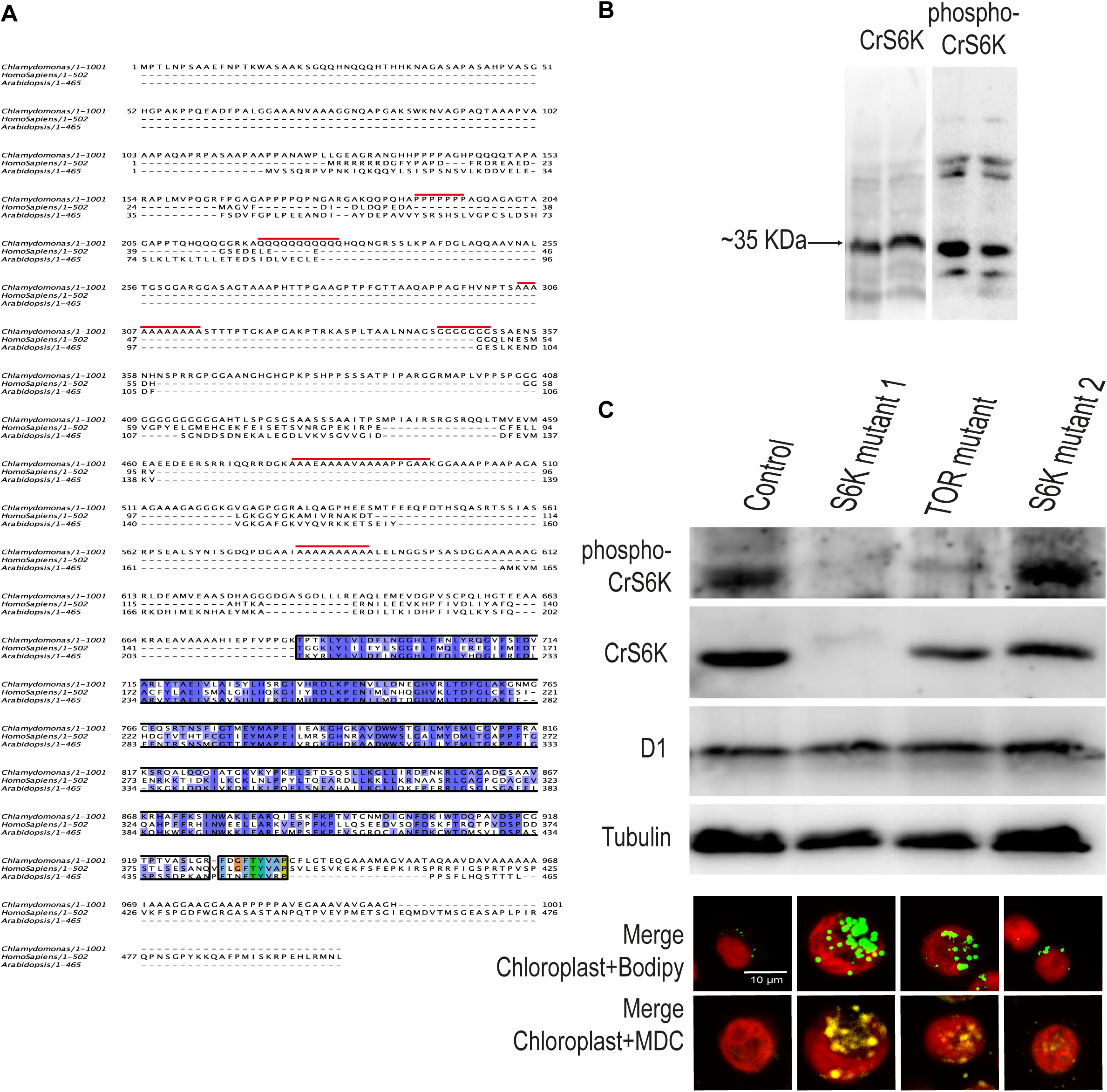
Prediction and identification of a putative CrS6K. **(A)** Multiple sequence alignment was performed across the annotated gene using MUSCLE of *C.reinhardtii, Arabidopsis* and Human S6K1 protein sequences. Thr residue in green at the extreme C-ter represents the conserved residue phosphorylated by TOR kinase within the TOR-motif (shown in box with other conserved residues in color). **(B)** The estimated size of the protein product from the blue region is ~35kDa, which is detectable in *C. reinhardtii* whole cell-extracts from log phase cells (grown in continuous light) using mammalian anti-S6K1 antibodies (western blot lanes from 10% SDS-PAGE gel shown in duplicate). **(C)** Western blot analyses of S6K mutant1, TOR mutant and S6K mutant 2 using the anti-phospho-S6K1, anti-S6K1, anti-D1 and anti-tubulin antibodies. Moreover, BODIPY dye staining of the mutants also shows lipid accumulation phenotype. Autophagosomes were stained using Monodansylcadaverine (MDC) and Chloroplast were observed using autofluorescence.

A western blot analysis using this antibody showed a prominent band at molecular weight of ~35 KDa suggesting that the band corresponds to the *C.reinhardtii* S6K1 protein (Fig.1B). In fact, we did not observe any cross-reactive band at the annotated molecular weight of 100 KDa. We also attempted to detect the phosphorylation status of Thr residue using a specific antibody against the Thr(P)-389 peptide of human pS6K1 (Cell Signaling Technology Cat. No.- 9234S). A similar phospho-specific Ab for the same epitope (Thr(P)-389) has been successfully tested in *Arabidopsis* (Cell Signaling Technology Cat no - 9205S) (Xiong *et al.*, 2012) and attempted previously in *C.reinhardtii* (Couso *et al*., 2016). We note that the human TOR phosphorylating motif has more similarity with that of *C.reinhardtii* (with only 1 change out of 9 residues) than *Arabidopsis* (3 changes out of 9 residues) as depicted (Fig. 1A). Even overexposed protein western blot against this antibody revealed phosphorylation of Thr(P) in S6K1 protein prominently at ~35 KDa (Fig. 1B). Other faint non-specific bands were common in both S6K1 and pS6K1-Ab westerns (Fig. 1B).

To further strengthen our claim on the molecular identity of CrS6K, we analysed the same in recently available S6K and TOR insertional mutants of *C.reinhardtii* (X. Li *et al.*, 2016) (obtained from https://www.chlamylibrary.org). We performed western blot analysis for bulk and pCrS6K in these mutant strains and observed that the 35 KDa band was absent in the S6K mutant 1 (LMJ.RY0402.091478) but not in S6K mutant 2 (LMJ.RY0402.209448). While characterizing the insertional mutants, it was noted earlier that the insertion of a cassette can lead to complete absence of the protein band as observed in lcs2 mutant (Fig. 7C in X. Li *et al.*, 2016). Moreover, the pCrS6K signal was also missing in the S6K mutant 1 while the level was substantially reduced in TOR mutant as compared to control wild type strain as well as unstable S6K mutant 2 (Fig. 1C). The absence of bulk CrS6K band in the S6K mutant 1 strongly suggested that the 35 KDa band indeed corresponds to *C.reinhardtii* S6K1 protein as argued by us on the basis of bioinformatics analysis. Moreover, the decrease in pCrS6K level associated with TOR mutant (LMJ.RY0402.203031) reinforced that the pCrS6K is an authentic functional readout of the endogenous TOR kinase activity. Additionally, S6K mutant 1 and TOR mutant exhibited autophagosomal structures constitutively, presumably due to high endogenous autophagy, consistent with their reduced TOR kinase activity and concomitant high lipid-body accumulation evident in these cells (Fig. 1C). We note that as shown in the X. Li *et al.*, 2016 study, some of the insertional mutants exhibit instability, as exemplified by S6K mutant 2 in the current study, which failed to show the expected mutant phenotype. Thus, based on S6K mutant 1 and TOR mutant data described here, for the first time, we argue that 35 kDa molecular species described here corresponds to *C.reinhardtii* CrS6K. We assayed CrS6K phosphorylation status as activity readout of endogenous TOR-kinase, which we probed to assess the metabolic plasticity in the cells.

### TOR kinase activity status (pCrS6K/CrS6K) is modulated by cellular metabolic states: comparison across phototrophy, mixotrophy and heterotrophy

We monitored TOR kinase activity in different growth conditions: Phototrophy (in light versus dark), mixotrophy and heterotrophy. We compared the same in N+ (Nitrogen replete) and N− (Nitrogen deficiency) conditions. Nutrient starvation is known to significantly impact TOR kinase activity (Gonzalez *et al.*, 2017; Loewith *et al.*, 2011). It has been well documented that autophagy is up regulated during nitrogen starvation and rapamycin treatment of *C.reinhardtii* cells (Perez-Perez, Couso, & Crespo, 2017; Perez-Perez, Couso, Heredia-Martinez*, et al.*, 2017). However, it is not known how TOR kinase activity is regulated in these metabolic states. Therefore, to establish TOR kinase activity status in *C.reinhardtii*, we monitored the phospho-CrS6K (pCrS6K) levels at different time points of nitrogen starvation in light versus dark conditions during autotrophy.

As can be seen in Fig. 2A, N-starvation, specifically in light, leads to a significant drop in bulk CrS6K level after 24h of incubation. The same effect was not observed in dark, reflecting the requirement of light signal in facilitating the N-starvation associated protein loss in algal cells. In dark, bulk CrS6K level is well maintained during N-starvation. Interestingly, even in early time-point of N-starvation (12h-24h) during light when the bulk CrS6K is maintained, that of pCrS6K level tends to drop as compared to the same time-points in N+ conditions (compare the ratios of pCrS6K to CrS6K at 12h & 24h in N+ with N− in Fig. 2A). pCrS6K/CrS6K level in N− cells drops to one third of N+ by 24h which becomes undetectably low after 24h-36h of N-starvation in light. In the same time-course, N-starvation response in dark shows no measurable change in pCrS6K level: the ratio of pCrS6K to CrS6K remains unchanged in N+ versus N− conditions in dark (compare the ratio values in Fig. 2A). These results show that TOR kinase activity drops during N-starvation specifically in light. Interestingly, bulk CrS6K level is well maintained during the time-course of N-starvation under mixotrophic conditions, both in light as well as dark conditions. But pCrS6K levels are significantly lowered in N-starvation under mixotrophy in both light and dark (i.e. heterotrophy) conditions (Fig. 2A), suggesting that TOR kinase activity loss during N-starvation does not depend on light signals during mixotrophic growth. The ratio of pCrS6K to S6K is similarly low (very low to detect) in both light and dark in these conditions during N-starvation (Fig. 2A).

**Figure 2.**
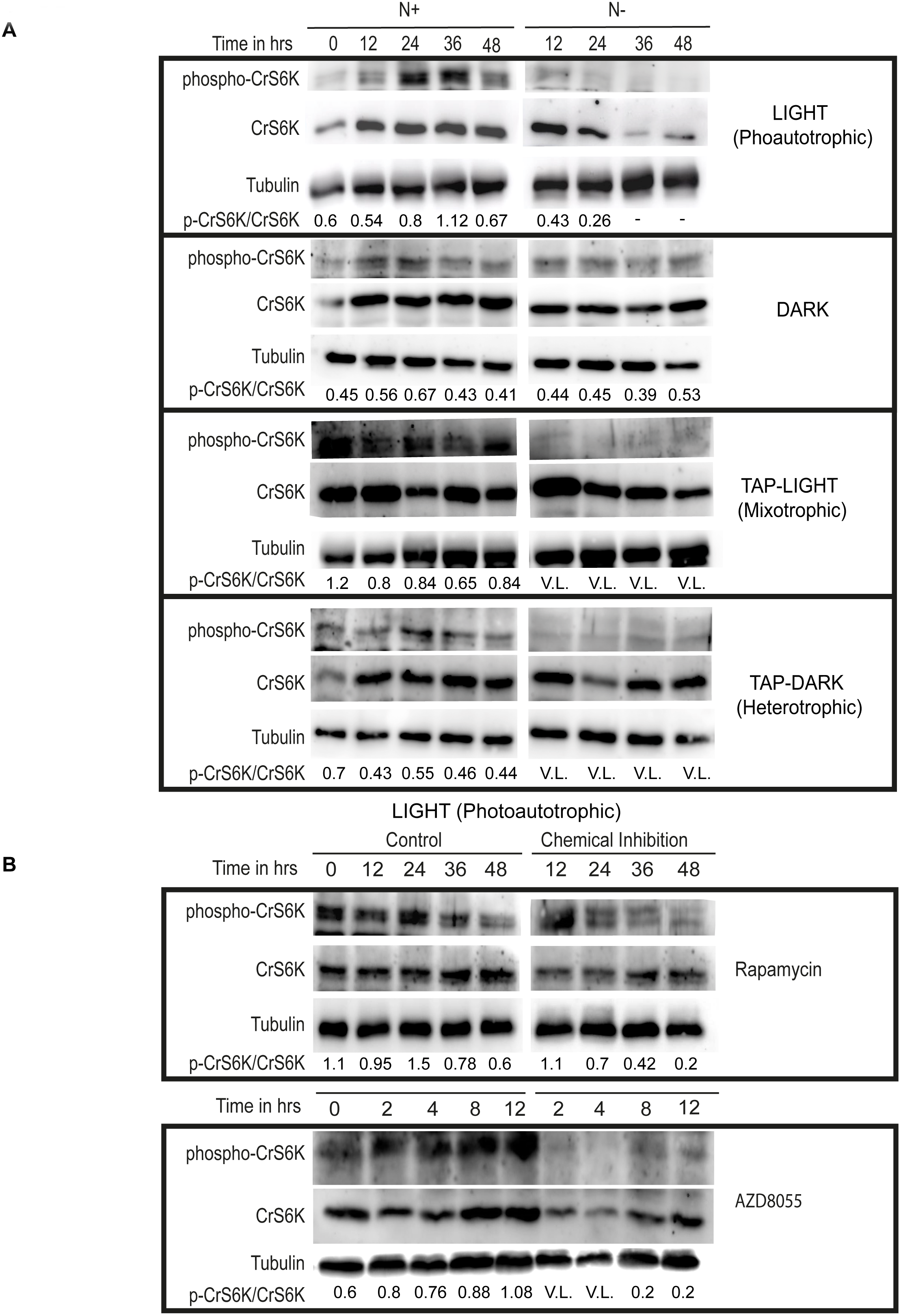
Regulation of putative CrS6K under various metabolic states. **(A)** *C. reinhardtii* cells from log-phase culture (photoautrophic) were shifted to different growth conditions, as specified (see Methods) in N-replete (N+) and N-deplete (N−) states. Samples were collected at different time-points of incubation, lysed, total protein (50 μg) was extracted and resolved on a 10% SDS-PAGE gel followed by western blotting using mammalian anti-p(T389)-S6K1 or anti-S6K1-Ab. **(B)** *C. reinhardtii* cells from log-phase culture were incubated with mTOR kinase inhibitors: Rapamycin (1 μM) or AZD8055 (1 μM) for different time points in photoautrophic conditions, followed by western blot analysis of total proteins using mammalian anti-p(T389)-S6K1 or anti-S6K1-Ab. pCrS6K/CrS6K ratios calculated using ImageJ intensity analysis where ‘-‘ indicates that the ratio cannot be calculated while V.L. signifies that the ratio is very low. Ratios were calculated as an average of 3 blots and normalized by Tubulin bands.

TOR-kinase inhibition, as reflected by the drop in pCrS6K/CrS6K ratio, is expected to enhance cellular autophagy as the two responses are known to act reciprocally (Jung *et al.*, 2010). In order to verify whether the drop in TOR kinase activity as revealed by decrease in pCrS6K level during N-starvation, specifically in light during autotrophy (but not in dark), also shows reciprocal change in autophagy, we assayed ATG8 protein level change, a mark of autophagy induction. Interestingly, western blot analyses revealed several fold enhancement of ATG8 during N-starvation in light as compared to that in dark (Supplementary Fig. 1A and Fig. 1D). Moreover, much higher chloroautophagy (fragmentation of chloroplast morphology, monitored by chlorophyll autofluorescence) was evident in light versus dark conditions. All these results put together strongly suggest that pCrS6K/CrS6K in the current study is a bona fide readout of TOR kinase activity status and therefore reflects a measure of cellular anabolic state, which we investigated further in the context of altered carbon metabolism in the cell.

We verified whether the chemical inhibition of TOR kinase via rapamycin or AZD8055 treatment results in expected reduction of endogenous TOR kinase activity, as measured by pCrS6K/CrS6K ratio analyses. Rapamycin, a specific inhibitor of TOR kinase has previously been used in *C.reinhardtii* to study autophagy induction and TOR kinase regulation of autophagy and oxidative stress (Diaz-Troya *et al.*, 2011; Perez-Martin *et al.*, 2014; Perez-Perez, Couso, & Crespo, 2017). AZD8055, a specific TOR kinase domain inhibitor (Benjamin *et al.*, 2011) has also been recently reported to bring about lipid accumulation in *C.reinhardtii* (Imamura *et al.*, 2016). When we treated *C.reinhardtii* cells with rapamycin or AZD8055, followed by western blot analyses, we observed that both rapamycin (1 μM) and AZD8055 (1 μM) resulted in the drop of pCrS6K/S6K. Rapamycin inhibition was evident after 24h treatment while AZD inhibition was set in by as early as 2h post treatment (see the ratios in Fig. 2B). These results showed that *C.reinhardtii* TOR kinase activity is well monitored by pCrS6K to CrS6K ratio measurement and that the kinase is efficiently inhibited by the inhibitor treatment, all pointing towards the robustness of the assay system described here in *C.reinhardtii* system for the first time.

### TOR kinase activity is negatively modulated by glucose: effects of atrazine and glucose on pCrS6K/CrS6K levels

Results described above uncovered light dependent attenuation of TOR kinase activity during autotrophy (Fig. 2A). In order to probe this effect further mechanistically, we performed the following experiment. We inhibited PSII electron transfer by treating the cells with atrazine (0.1 mM) (Moreland, 1980), and followed pCrS6K/S6K levels as a function of time which revealed that the ratio remains unchanged in N+ versus N− conditions in the entire time-course when PSII was inhibited (Fig. 3A). Quantification of the ratio revealed remarkable stability of TOR kinase activity in autotrophy even in light when PSII was inhibited, thereby phenocopying the effect of autotrophy in dark, a result that starkly contrasts with light dependent attenuation of TOR kinase activity when PSII was active (i.e. normal photosynthesis)(Fig. 2A).

**Figure 3.**
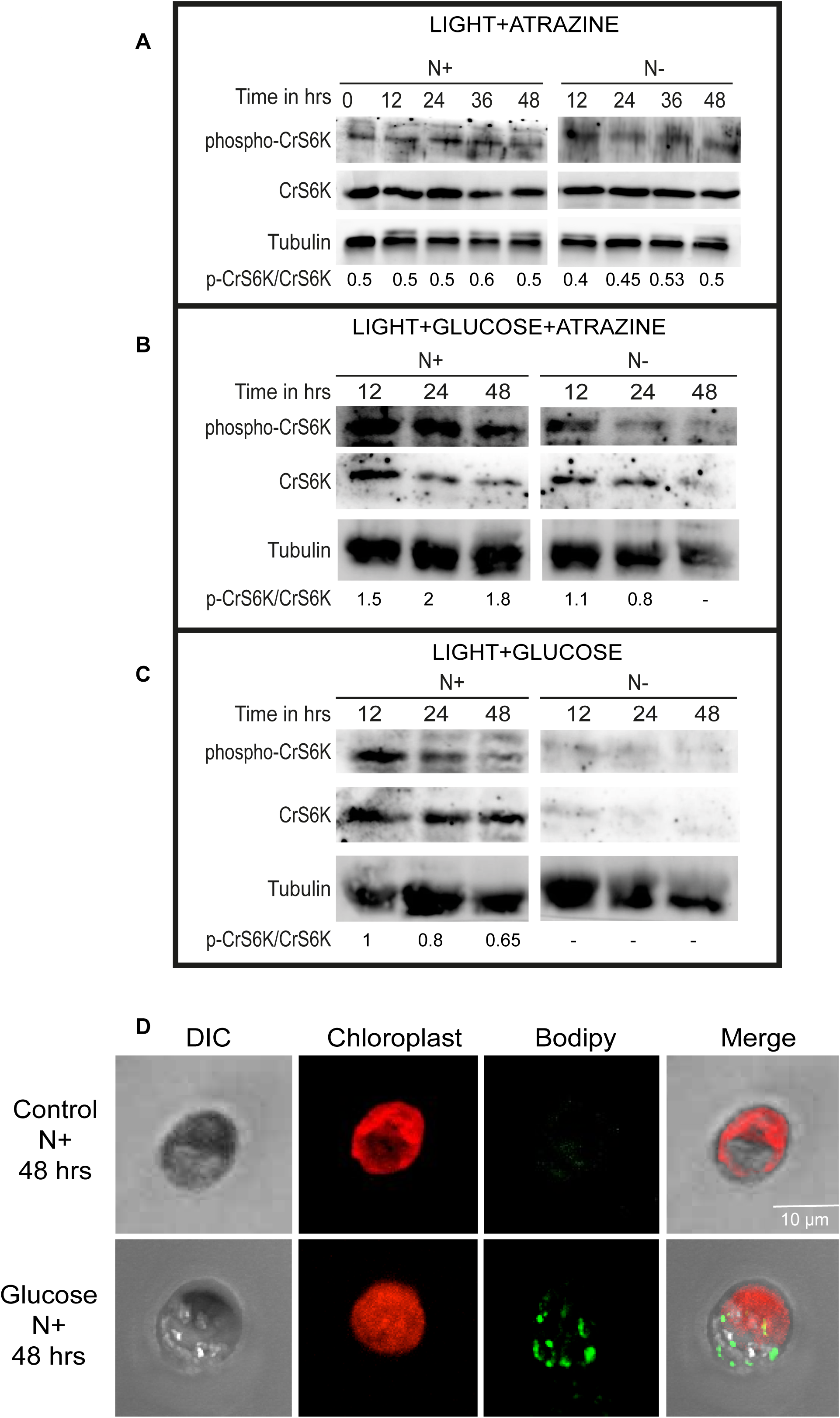
Effect of Atrazine treatment and glucose addition. *C. reinhardtii* cells from log-phase culture (photoautrophic) in N-replete or N-deplete states were incubated with **(A)** Atrazine (0.1 mM) or **(C)** D-Glucose (100 μM) or **(B)** both, followed by western blot analyses of samples retrieved at different time-points of treatments. **(D)** *C. reinhardtii* cells from log-phase culture (photoautrophic) in N-replete (Control) or the same with D-Glucose (100 μM) were collected after 48 h, stained with BODIPY and imaged using Zeiss Confocal 510 microscope. Lipid bodies were visualized following BODIPY staining and imaged for fluorescence at 488 nm. Chloroplast morphology was imaged by autofluorescence at 600 nm while DIC imaging revealed the shape of the cell.

Based on this result, we surmised that photosynthetic output; perhaps organic carbon (or its metabolic product) could be responsible for TOR kinase inhibition during autotrophy, specifically in light. Photosynthesis results in the fixation of carbon and formation of metabolically important sugars, especially sucrose and glucose (L. Li *et al.*, 2016) that have also been shown to act as signaling molecules. Especially, glucose is known to activate TOR kinase in *Arabidopsis* (Xiong *et al.*, 2013). We tested the same by supplementing the light grown and atrazine treated cultures with exogenous glucose (100 μM). Surprisingly, we observed a measurable drop in TOR kinase activity specifically in N− by 12h of glucose addition as compared to N+ cells (Fig. 3B). The same effect was accentuated further at later time points (i.e. 24-48h), suggesting that photosynthetic output, organic carbon, might impose some form of negative effect on TOR kinase activity in the cell, perhaps via a feedback regulation, specifically during N starvation (see Discussion). We tested the effect of glucose addition more directly in a control experiment involving no atrazine treatment. Even in this control, addition of glucose led to a dramatic loss of bulk CrS6K protein, specifically in N− condition (Fig. 3C). Concomitantly, pCrS6K levels also dropped too low to be detected in glucose supplemented N− cells, implying that addition of glucose accentuated the N starvation effect (loss of TOR kinase) in these cells (compare N− in Fig. 2A with 3C), suggesting that excess cellular carbon (resulting from glucose supplementation) mimics N− state in the cells perhaps via unbalancing the cellular C/N pool ratio (see Discussion). In fact, glucose induced N-starvation effect (TOR kinase attenuation and lipid accumulation) became evident even in N+ cells following glucose supplementation (Fig. 3C and Fig. 3D). pCrS6K/CrS6K ratio showed a steady decline as a function of time even in N+ cells following glucose addition (compare the ratios in Fig. 3C). Concomitantly, cells showed lipid accumulation following glucose supplementation, even in the absence of N starvation (N+ state) (Fig. 3D).

### Glucose uptake by *C.reinhardtii* cells leads to induction of autophagy, rise in ROS levels and changes in TOR kinase activity status

Earlier reports suggested that *C.reinhardtii* cells lack glucose uptake mechanism and that they cannot grow heterotrophically (dark) in the presence of glucose as the sole carbon source (Doebbe *et al.*, 2007; Karpagam *et al.*, 2015; Sager *et al.*, 1953). However, the uptake of glucose in *C.reinhardtii* cells in light versus dark has not been carefully studied so far. It is also important to note here that some *Chlamydomonas* species are known to utilize glucose efficiently to support their growth both hetero- and mixotrophically (Bennett *et al.*, 1972; Karpagam *et al.*, 2015). In order to rationalize glucose addition effects observed in the current study, we probed the uptake of glucose by *C.reinhardtii* cells in our experimental conditions.

To test whether *C.reinhardtii* cells exhibit glucose uptake in light *versus* dark, we used a fluorescently tagged non-hydrolysable analogue of deoxyglucose i.e 2-NBDG (NBD Glucose) that fluoresces only upon entering into live cells, but cannot be hydrolyzed further following its conversion to glucose-6-phosphate (Yamada *et al.*, 2007; Yoshioka *et al.*, 1996). NBDG fluorescence has been used in several cellular model systems as a robust indicator of glucose uptake (Etxeberria *et al.*, 2005; Zou *et al.*, 2005). We used the same to quantitatively assess glucose uptake by *C.reinhardtii* cells. We incubated the cells with 2-NBDG (0.1 mM) and measured fluorescence increase as a function of time in light versus dark incubation using microplate fluorescence reader (Tecan Infinite M1000). Interestingly, the uptake of 2-NBDG by cells was seen only in light, but not in dark (Fig. 4A). We observed a slow linear increase in the NBDG fluorescence, which tends to saturate by about 8-10h in light. We corroborated glucose uptake result by performing glucose concentration dependent growth assay where we note a marginal, but measurable increase in cell density that was glucose concentration dependent. In fact, the growth level showed an optimum at certain glucose concentration (80 μM) in these conditions (Fig. 4B and Fig. 4C). These results support the claim that addition of glucose leads to discernible level of growth in *C.reinhardtii* culture specifically in mixotrophic conditions (light+glucose). The same in dark leads to no glucose uptake (Fig. 4A) and consequently no growth in the culture where prolonged incubation leads to cell-death (data not shown).

**Figure 4.**
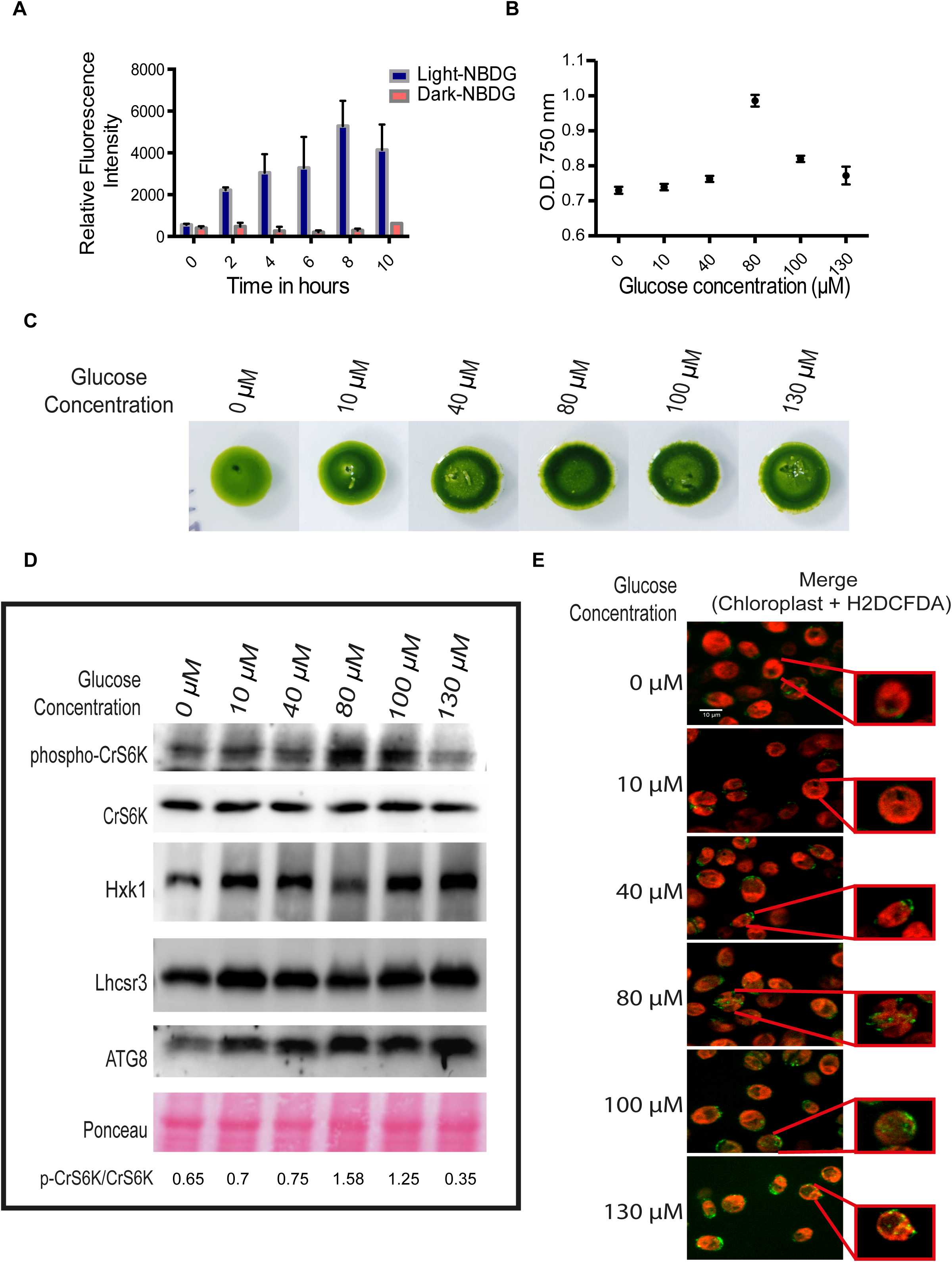
Glucose uptake and utilization by *C. reinhardtii* cells. **(A)** *C. reinhardtii* cells from log-phase culture (photoautrophic) were incubated with 2-NBDG (0.1 mM) (fluorescently tagged deoxy-glucose) in light or dark, followed by fluorescence measurement (Excitation at 400 nm and emission at 550 nm) at 0, 2, 4, 6, 8 and 10 h of incubation using Tecan Infinite M1000 fluorescence plate reader. *C.reinhardtii* cells from log-phase culture (photoautrophic) were incubated at different concentrations of glucose (0, 10, 40, 80, 100 and 130 μM), followed by cell density analyses after 48 h of growth (absorbance at 750 nm) **(B)**. Colony images on TP agar plates after 5 days of growth **(C). (D)** Samples in 4B were extracted for total protein, resolved on 10% SDS-PAGE gel, followed by western blotting using mammalian anti-p(T389)-S6K1-Ab or anti-S6K1-ab or anti-ATG8 or *C.reinhardtii* anti-Hxk1 or anti-Lhcsr3 Ab’s. **(E)** Samples from experiment described in 4B were imaged live for ROS using H2DCFDA (10 μM) staining or chloroplast imaging using autofluorescence (excitation at 500 nm, emission at 600 nm).

To investigate whether the glucose uptaken by the cells also participates in signal transduction, we performed a western blot analysis of cells for Hexokinase1 (Hxk1) and pCrS6K levels in cultures treated with different concentrations of glucose. Just as growth (Fig. 4B and Fig. 4C), pCrS6K level also showed an optimum response at a particular concentration of glucose (80 μM) (compare the pCrS6K/CrS6K ratios in Fig. 4D). Hxk1 is known to act as a direct sensor and integrator of glucose signals in plants (Cho *et al.*, 2006; Moore *et al.*, 2003). However, the same in *C.reinhardtii* has not been well characterized. Interestingly, Hxk1 level was elevated following glucose addition, suggesting that cells were sensing the organic carbon in the medium. It has been suggested that excess glucose can result in autophagy induction and increase in ROS (Reactive Oxygen Species) levels (Moruno *et al.*, 2012; Yu *et al.*, 2006), which was also evident in our experiment by the rise in autophagy as monitored by ATG8 levels (Fig. 4D). Interestingly, glucose addition as a carbon source resulted in systematic increase in intracellular ROS levels detected by the ROS-specific dye H2DCFDA positivity (Fig. 4E). In spite of rise in ROS levels and the concomitant induction of autophagy marker (ATG8), cells showed normal chloroplast morphology. Photosynthetic function as monitored by a Non-Photochemical Quenching (NPQ) marker (Lhcsr3 protein) showed no discernible changes during the entire range of glucose addition (Fig. 4D). Consistent with autophagy induction, cells exhibited lipid accumulation at different glucose concentrations (Supplementary Fig. 2). Thus, all these data put together, strongly justify our hypothesis that *C.reinhardtii* cells exhibit glucose uptake, respond to glucose signals and evoke various physiological responses such as autophagy induction, lipid accumulation, along with the concomitant rise in ROS levels, specifically in light. In order to probe the mechanistic relationship between ROS, autophagy and TOR kinase changes associated with glucose uptake by the cells in light, we tested whether ROS itself acts as an upstream regulator of autophagy/TOR kinase activity changes. Incubation of cells with increasing exogenous level of H_2_O_2_ (ROS inducer) in the medium resulted in concentration dependent drop in TOR kinase activity and a reciprocal rise in ATG8 level in the cells (Supplementary Fig. 3). Glucose concentration dependent rise followed by a drop in TOR kinase activity status, leading to an optimum TOR kinase activity at a particular glucose level (80 μM) (Fig. 4D) suggests that glucose mediated change in TOR kinase activity follows a bi-phasic response: first phase where TOR kinase activity rise is perhaps ROS independent and glucose (or glucose derived metabolites) dependent (see Discussion) followed by the second phase where increase in ROS leads to attenuation of TOR kinase activity at high glucose level (130 μM). We discuss this model further integrating other results described here (Fig. 5).

**Figure 5.**
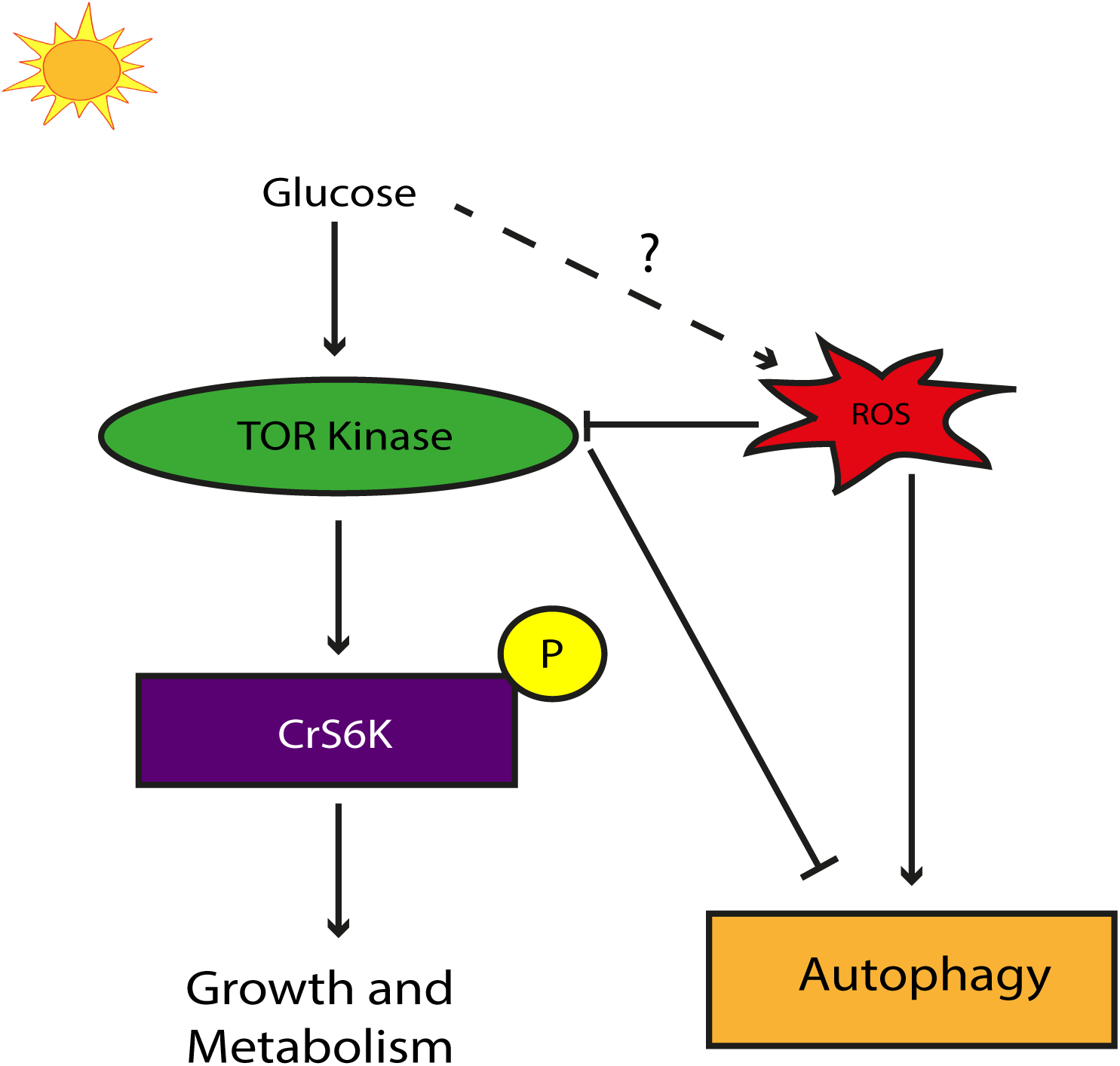
A model suggesting that TOR kinase receives positive inputs from glucose whereas at the same is under the negative regulation by ROS giving rise to a bi-phasic modulation of TOR kinase in light grown autotrophic cultures.

## DISCUSSION

One of the most well characterized downstream targets of TOR kinase is the Ribosomal protein S6 Kinase (Sabatini, 2017; Saxton *et al.*, 2017; Schalm *et al.*, 2002). TOR kinase phosphorylates its target, S6 Kinase, at the Thr-389 in the mammalian S6K. The N-terminus of S6K has a TOR signaling (TOS) motif where Raptor and TOR kinase bind to phosphorylate the linker region in the C-terminus at Thr-389 residue in mammals (Schalm *et al.*, 2002). The TOS motif is conserved in animals and yeast. However, plants have evolved a different TOS motif suggesting that they have a different mechanism of binding and activation of S6K (Yaguchi *et al.*, 2018). Thus, although the C-terminal linker region has the conserved Thr residue that acts as a phosphorylation target of TOR kinase, the N-terminal region is not conserved. This is what we observe in our MUSCLE alignment where, the C-terminal region of *C.reinhardtii* is highly conserved (as seen in blue in Fig. 1A), while the N-terminal region was non-conserved.

Moreover, our western blot analyses of *C.reinhardtii* S6K protein (CrS6K) shows a single band at ~35 KDa, while no band is observed at the predicted molecular weight of 100 KDa (Fig. 1B). Thus, it is likely that the N-terminal non-conserved region is spliced off leading to the mature product of putative molecular weight of *C.reinhardtii* S6K is 35 KDa. The Thr(P)-389 S6K specific mammalian antibody also shows a band at the corresponding 35 KDa position indicating that it could be used as readout for the endogenous activity of TOR kinase.

We also used the recently made available S6K and TOR insertional mutants (X. Li *et al.*, 2016) to assess both the level of pCrS6K as a measure of TOR kinase activity and also to confirm the molecular weight of CrS6K. These mutants are generated by random insertion of a paramomycin cassette followed by screening for insertions within specific genes. S6K mutant 1 shows complete absence of both the pCrS6K and CrS6K protein bands at 35 KDa whereas the TOR mutant shows reduced level of pCrS6K, while the level of bulk CrS6K is maintained. Thus, these mutant studies reinforce our claim that the cross-reactive bands in western blot for pCrS6K and bulk CrS6K were authentic with respect to their putative molecular weight of ~35 KDa. However, due to certain insertions being unstable, partial or in the opposite orientation within the coding region, some mutants do not show the complete loss of protein band as observed in S6K mutant 2. Moreover, the S6K mutant 1 and TOR mutant show lipid accumulation (BODIPY staining) and autophagosomal structures (Monodansylcadaverine staining) (Fig. 1C) constitutively, thereby phenocopying *C.reinhardtii* cells under nutrient starvation, indicating that the basal state of these mutants is similar to starved state in the presence of TAP media (with normal carbon and nitrogen supplement).

Having developed an endogenous assay to monitor TOR kinase activity in *C.reinhardtii*, we further probed the changes in TOR kinase activity under different physiological conditions. Previous studies with TOR kinase inhibition have shown that it is important for biomass accumulation, cell cycle progression and maintenance of carbon metabolism in *C.reinhardtii* (Juppner *et al.*, 2018; Kleessen *et al.*, 2015). Our results indicate that the TOR kinase activity in N+ state under photoautotrophy increases as a function of time up to 24h and then decreases (Compare pS6K to S6K ratios in Fig. 2A). Thus, as overall cell density or biomass increases, pCrS6K levels also increase and subsequently fall due to higher cell density or cell growth perhaps due to a feedback signal that regulates TOR kinase activity. The mechanism of feedback regulation of TOR kinase by cell density or cell cycle related effects need to be probed further. Under N starvation, however, there is a decrease in pCrS6K and bulk CrS6K as a function of time, which suggests that starvation acts as a catabolic signal and results in inhibition of TOR kinase and also causes catabolic breakdown of bulk CrS6K proteins. Previous studies have shown that the target of S6K, the ribosomal protein S6 (RPS6), undergoes degradation upon N starvation which is regulated by autophagy in *C.reinhardtii* (Couso *et al.*, 2017). Thus, it is possible that the degradation of bulk CrS6K is also under autophagy regulation.

In photosynthetic organisms, unlike in mammals and yeast, N starvation involves two opposing signals: the catabolic N starvation signal and the anabolic light/photosynthetic carbon fixation signal. To test whether the anabolic signal is also sensed by TOR kinase, we also performed the N starvation assay in the absence of the anabolic signal i.e. in the dark and monitored TOR kinase activity. Surprisingly, we observe that in dark, N starvation does NOT result in a decrease in pCrS6K level (Fig. 2A) indicating that, in the absence of any anabolic signal (light/photosynthetic carbon fixation), the catabolic signal (Nitrogen starvation alone) is not perceived by TOR kinase. To test the role of light alone in regulating TOR kinase activity, we inhibited photosynthesis by atrazine. This would prevent any carbon fixation but the effect of light alone on TOR kinase activity remains unaffected. Upon atrazine treatment, we observe that the pCrS6K levels do not change under N starvation (Fig. 3A), a result similar to N starvation in the dark, indicating that light itself may not act as a signal to regulate TOR kinase activity during N starvation.

We further tested the role of organic carbon (photosynthetic output) in regulating TOR kinase activity under starvation. When glucose was provided as the photo-assimilate carbon source under N starvation during photosynthesis inhibition (atrazine treatment), the pCrS6K levels reduce indicating that the carbon source i.e. an anabolic signal along with N starvation can result in reduction of TOR kinase activity, thereby suggesting that additional carbon in the cells attenuate TOR kinase activity, a result counterintuitive to that reported in mammalian system (Fig. 3B). Previous studies have shown that TOR kinase inhibition can disrupt the Carbon/Nitrogen (C/N) balance (Juppner *et al.*, 2018), while our results strongly indicate that disrupting the C/N balance can in turn result in inhibition of TOR kinase activity suggesting that the central metabolic sensor TOR kinase can sense as well as respond to metabolic signals in a feedback regulatory circuit. Interestingly, when we provide exogenous glucose to photoautotrophic cells under N starvation, the starvation response is accentuated further by dramatic attenuation of TOR kinase activity as monitored by pCrS6K levels (Fig. 3C). Moreover, even in N+ state, glucose addition results in measurable attenuation of TOR kinase activity, autophagy induction accompanied by lipid accumulation, hallmark signatures of starvation in *C.reinhardtii*. All these surprising results point towards the effect of added glucose that tip the cellular C/N balance, thereby creating “starvation like” phenotype, leading to a drop in TOR kinase activity. These results are contradictory to the established notion that glucose acts as an upstream positive regulator of TOR kinase activation.

Moreover, we note that no plasma membrane hexose transporters are reported in *C.reinhardtii* till date. In fact, it is largely believed that *C.reinhardtii* fail to uptake and grow on glucose as the sole carbon source in heterotrophic (dark) conditions (Doebbe *et al.*, 2007; Sager *et al.*, 1953). However, our results suggest that glucose is sensed and utilized in light and regulates TOR kinase activity (Fig. 4D). Thus, we probed this aspect further and discovered that the uptake of glucose, monitored by the fluorescently tagged 2-NBDG (NB-deoxyglucose), ensues only in the presence of light and not in dark (Fig. 4A) in *C.reinhardtii* cells. Moreover, growth on plates and absorbance assays with different concentrations of glucose in light suggests that there is an optimum concentration of glucose (80 μM) that results in maximum growth (Fig. 4B and Fig. 4C). To further test how TOR kinase activity is regulated under different glucose concentrations and also to explain the TOR kinase inhibition by excess glucose, we monitored pCrS6K levels across varying levels of glucose supplementation. Interestingly, we observe a bi-phasic response, where TOR kinase activity is positively correlated with glucose increase (in low glucose regime), while higher glucose concentration inhibits TOR kinase (Fig. 4D). Moreover, ATG8, which should follow an opposite trend to that of pCrS6K, shows rise at all levels of increasing glucose concentration. It has been previously reported in mammalian cells and yeast that a hyperglycemic (excess glucose) state results in generation of ROS species (Moruno *et al.*, 2012; Yu *et al.*, 2006). These ROS species can act as oxidative stress in *C.reinhardtii* and result in induction of autophagy (Perez-Perez *et al.*, 2012; Pérez-Pérez *et al.*, 2017). To test this hypotheses, we monitored the level of ROS by H2DCFDA dye staining and observed an increase in ROS with increasing glucose concentration (Fig. 4E) indicating that oxidative stress can cause induction of ATG8 as observed in Fig. 4D. Oxidative stress alone generated by treating the cells with sub-lethal levels of H_2_O_2_ can cause reduction in pCrS6K as seen in Supplementary Fig. 3, hinting at a plausible mechanism for inhibition of TOR kinase activity at higher glucose concentrations (130 μM).

Previous studies have shown that glucose activation of TOR kinase is independent of light and phytohormones (Li *et al.*, 2017), but the mechanism of activation at lower glucose concentrations (10-80 μM) is far from clear. Other important metabolic sensors such as SnRK’s (Snf Related protein Kinases) that are homologs of AMPK, but may sense glucose/glucose-6-phosphate levels in plants along with ATP/AMP levels (sensed by AMPK) have been implicated to directly interact with Raptor and TOR kinase to regulate its kinase activity (Li *et al.*, 2017; Robaglia *et al.*, 2012; Soto-Burgos *et al.*, 2017; S. Wang *et al.*, 2015). However, further studies need to be carried out to carefully understand the signaling mechanism of glucose sensing by SnRK’s as well as TOR kinase, and the underlying co-regulation, if any. Recently available insertional mutants of different SnRK proteins and TOR kinase in C.reinhardtii may greatly help to discern these processes involved in glucose sensing and signaling.

## MATERIALS AND METHODS

### Strains, media and growth conditions

*C.reinhardtii* WT CC-125 was obtained from the *C.reinhardtii* Resource Center (https://www.chlamycollection.org). *C.reinhardtii* insertional mutants S6K mutant 1 (LMJ.RY0402.091478), S6K mutant 2 (LMJ.RY0402.209448) and TOR mutant (LMJ.RY0402.203031) are a set of newly available insertional mutants which were obtained from the *Chlamydomonas* Library Project (CLiP) (https://www.chlamylibrary.org). *C.reinhardtii* WT cells CC-125 were grown under continuous illumination at 25^°^C in Tris Phosphate (TP) for photoautotrophy, Tris Acetate Phosphate (TAP) in light for mixotrophy and TAP in dark for Heterotrophy; while the mutants were grown in TAP media as described (E. Harris *et al.*, 1989). For nitrogen deprivation, photoautotrophically grown cells (10^6^ cells/ml) were pelleted and resuspended in TP N-media. Cells in exponential phase (10^6^ cells/ml) were treated with atrazine (Sigma, 49085), or rapamycin (MCE, HY-10219), or AZD8055 (Axon medchem, Axon 1561), or NBDG (Cayman Chemical, 11046 – 1 mg), or CM-H2DCFDA (ThermoFischer, C6827) as described in the experiments. Media was also supplemented with glucose (Merck Millipore, CAS 50-99-7) for some experiments. Additionally, glucose supplemented TP Agar (1% Bacto Agar) plates were used to grow cells in light in presence of glucose at various concentrations. Optical density of cells was measured as absorbance at 750 nm to get and estimate of cell density and number as shown previously (Burgess *et al.*, 2016).

### Protein preparation and immunoblot analysis

Total proteins from *C.reinhardtii*, (corresponding to 1 μg of chlorophylls, used as input normalisation), were extracted by pelleting exponentially grown cells (800 *g* for 10 min) and resuspending the pellet with Laemlli buffer (1M Tris-HCl pH 6.8; 10% SDS (w/v); 5% β-mercaptoethanol; 0.02% Bromophenol blue) and subsequently denaturing at 100°C (Heating block) for 20 min. The samples were cooled by keeping them on ice for 5 min. Cell debris was removed by centrifugation at 20,000 *g* for 15 min. The supernatant is used for protein western blotting. Samples were normalized with chlorophyll content and proteins were separated on 10% and 15% (for ATG8) SDS-PAGE and blotted for 2 hrs at 100 V by transferring on nitrocellulose membrane. Blots were blocked with blocking solution (PBS 1X, 0.2% w/v Tween, 5% powder milk) for 1h at room temperature (RT) with agitation. Blot was incubated in the primary antibody diluted in blocking solution for 1h at RT with agitation. The antibody solution was decanted and the blot was rinsed briefly twice, then washed 3 times for 10 min in blocking solution at RT with agitation. Blot was incubated in secondary antibody (anti-rabbit IgG Peroxidase conjugated; Sigma, A9169; 1:10,000 dilution in blocking buffer) for 1h at RT with agitation. The blot was washed 2 times for 10 min in blocking solution and once with PBS 1X solution for 10 min, then developed in developing buffer. Primary antibodies anti-Phospho p70 S6 Kinase Thr389 (Cell Signalling Technology (CST), 9234), anti-p70 S6 Kinase (CST, 9202), anti-Tubulin alpha chain (Agrisera, AS10 680), anti-D1 protein of PSII (Agrisera, AS05 084), anti-LHCSR3 (Agrisera, AS14 2766), anti-Hxk1 (Agrisera, AS16 4083, kindly provided by Joanna Porankiewicz-Asplund), anti-APG8A (Abcam, ab77003) were used at a dilution of 1:1000, 1:1000, 1:10,000, 1:10,000, 1:1000, 1:1000 and 1:1000 respectively. The blots were further developed using Chemiluminiscence (Biorad Clarity Max western ECL blotting substrates, 1705062).

### Immunofluorescence microscopy

For immunofluorescence staining, *C.reinhardtii* cells were immobilized on <u>Poly-L-lysine solution 0.1% (w/v) in H_2_O</u> (Sigma-Aldrich) coated slide, followed by fixation with 1.8% paraformaldehyde (Sigma-Aldrich) for 20 min. Slides were washed with 0.1 M glycine (Sigma-Aldrich) for 30 min and permeabilized using Triton X-100 (GE Healthcare) for 8 min. The cells were blocked using 2% BSA prepared in phosphate-buffered saline (PBS), incubated with anti-APG8A primary antibody (1:200) (Abcam, ab77003) followed by incubation with anti-rabbit secondary antibody tagged with Alexa Fluor 488 (Life technologies) (1:500). The cells were imaged in a Zeiss 510 Laser Scanning Confocal Microscope.

### MDC staining for in vivo labeling of autophagosomes

MDC (Monodansylcadaverine) staining of *C. reinhardtii* cells was performed as previously described (Contento *et al.*, 2005). Briefly, 1×10^6^ cells of nitrogen starved (N−) and non-starved (N+) cells were collected by centrifugation and re-suspended in of 100 μL TP media and stained by incubating with 50 μM MDC for 10 min at room temperature. The cells were then washed with PBS twice to remove excess dye and observed using fluorescence microscope.

### BODIPY staining of lipid bodies in C. reinhardtii

BODIPY 493/503 (4,4-difluoro-1, 3, 5, 7, 8-pentamethyl-4-bora-3a, 4a-diaza-s-indacene), a lipophilic fluorescent dye (Life technologies) was used for detecting lipid bodies in *C. reinhardtii* as previously described by Work et al., 2010. 1×10^6^ cells in a total volume of 100 μL were suspended in a final concentration of 5 μg/mL BODIPY using a stock concentration of 10 mg/mL BODIPY dissolved in dimethyl-sulfoxide. The cells were incubated in dark for 10 min and washed thrice in TP medium. Images were acquired using Zeiss 510 Laser Scanning Confocal Microscope. BODIPY dye with a 493 nm excitation and 503 nm emission wavelengths was detected using a 515/30 band-pass filter.

### NBDG uptake by *C.reinhardtii*

NBDG (2-deoxy-2-[(7-nitro-2,1,3-benzoxadiazol-4-yl)amino]-D-glucose), a fluorescently labeled deoxyglucose analog is used to monitor glucose uptake by living cells and tissues. 1 ml culture of exponentially grown *C.reinhardtii* cells were treated with fluorescently tagged 0.1 mM 2-NBDG and incubated in continuous light and dark on a rotating platform. Fluorescence was measured at various time points using a Tecan Infinite M1000 plate reader at excitation and emission wavelengths of 400 nm and 550 nm respectively. Relative fluorescence readings were measured and their average was plotted using Prism Software. Statistical analysis of standard deviation done using Prism software.

### ROS staining by CM-H2DCFDA

CM-H_2_DCFDA (ThermoFischer, C6827) is a cell permeable fluorescent probe 6-chloromethyl-2’,7’-dichlorodihydrofluorescein diacetate used to visualize ROS in *C.reinhardtii* cells. H_2_DCFDA staining was performed as described in (Vallentine *et al.*, 2014). Briefly, 10^6^ cells were incubated in 1 ml TP media containing 10 μM H_2_DCFDA (dissolved in DMSO) and incubated on a rotating platform in the dark for 20 min. Images were acquired using Zeiss 510 Laser Scanning Confocal Microscope. Chlorophyll auto-fluorescence was detected using a long-pass optical filter of 600 nm and H2DCFDA with and excitation wavelength of 492 nm and an emission wavelength of 525 nm was detected using a 500-530 nm band-pass optical filter.

## Acknowledgements

We thank Joanna Porankiewicz-Asplund for providing us with Hxk1 antibody. We acknowledge JC Bose fellowship grant (DST) [10X-217 to B.J.Rao]; and Department of Atomic Energy (Government of India) grant to Tata Institute of Fundamental Research (TIFR), Mumbai [12P0123].

## Supplementary Figures

**Supplementary Figure 1. Autophagy induction in light and dark.** (A) Log phase cells were transferred to N-replete (N+) and N-deplete (N−) media and incubated in light. Samples were collected at 12, 24, 36, 48 h and immunofluorescence using anti-ATG8 antibody was performed. Cells were imaged using the Zeiss 510 confocal microscope. (B) Above mentioned treatment was performed in light and dark and samples were collected for protein extraction. 50 μg of protein was run on a 15% SDS-PAGE gel followed by western blotting using the anti-ATG8 antibody. (C) Log phase cells were transferred to N-replete and N-deplete media and incubated in dark. Samples were collected at 12, 24, 36, 48 h and immunofluorescence using anti-ATG8 antibody was performed. Cells were imaged as mentioned before. (D) Log phase cells were transferred to N-replete and N-deplete media and incubated in light and treated with 0.1 mM Atrazine. Samples were collected at 12, 24, 36, 48 h and immunofluorescence using anti-ATG8 antibody was performed. Cells were imaged as mentioned before.

**Supplementary Figure 2: Lipid accumulation in different glucose concentrations.** *Chlamydomonas* cells in exponential growth were transferred to media containing following concentrations of glucose (0, 10, 40, 80, 100 and 130 μM). Cells were collected after 48 hrs and stained with BODIPY to assess lipid bodies. Live imaging using Zeiss Confocal 510 microscope was performed to analyze lipid bodies at 488 nm and autofluorescence of chloroplast at 600 nm.

**Supplementary Figure 3. Autophagy induction by H_2_O_2_.** (A) Log phase cells were treated with different concentrations of H_2_O_2_ and samples were collected 8 h post treatment. 50 μg of protein was run on a 15% SDS-PAGE gel followed by western blotting using the anti-ATG8, anti-pS6K, anti-S6K and *Chlamydomonas* anti-Hxk1 and anti-Tubulin antibodies. Dye staining using H2DCFDA to detect various species of ROS was performed. Cells were imaged using the Zeiss 510 confocal microscope.

Author Contributions S.U. and B.J.R. conceived and planned the research. S.U. performed the experiments with help from A.G., S.A. while S.U. and B.J.R. wrote the Manuscript.

